# AnnoTree: visualization and exploration of a functionally annotated microbial tree of life

**DOI:** 10.1101/463455

**Authors:** Kerrin Mendler, Han Chen, Donovan H. Parks, Laura A. Hug, Andrew C. Doxey

## Abstract

Bacterial genomics has revolutionized our understanding of the microbial tree of life; however, mapping and visualizing the distribution of functional traits across bacteria remains a challenge. Here, we introduce AnnoTree - an interactive, functionally annotated bacterial tree of life that integrates taxonomic, phylogenetic, and functional annotation data from nearly 24,000 bacterial genomes. AnnoTree enables visualization of millions of precomputed genome annotations across the bacterial phylogeny, thereby allowing users to explore gene distributions as well as patterns of gene gain and loss across bacteria. Using AnnoTree, we examined the phylogenomic distributions of 28,311 gene/protein families, and measured their phylogenetic conservation, patchiness, and lineage-specificity. Our analyses revealed widespread phylogenetic patchiness among bacterial gene families, reflecting the dynamic evolution of prokaryotic genomes. Genes involved in phage infection/defense, mobile elements, and antibiotic resistance dominated the list of most patchy traits, as well as numerous intriguing metabolic enzymes that appear to have undergone frequent horizontal transfer. We anticipate that AnnoTree will be a valuable resource for exploring gene histories across bacteria, and will act as a catalyst for biological and evolutionary hypothesis generation.

## Introduction

Important biological and evolutionary insights can be generated by exploring the presence/absence of genes and functional annotations across species phylogenies. These include identifying unexpected taxonomic occurrences (Venter et al. 2004), uncovering the evolutionary origin of genes (Demuth and Hahn 2009), and locating putative horizontal gene transfer (HGT) events (Andersson et al. 2006; Ravenhall et al. 2015). With the ongoing exponential increase in available genome sequences, including information from previously uncharacterized and uncultured lineages, online genomic repositories are becoming increasingly valuable collections of predicted genes and functional annotations. With this wealth of genomic data comes the opportunity for large-scale examinations of gene family distributions and evolutionary histories, but databases are not easily accessed, updated, or visualized.

A number of strategies exist for merging taxonomic and functional information to create annotated phylogenies. For instance, homologs of a gene family retrieved using BLAST (Camacho et al. 2009) or related methods can be manually mapped onto a custom species tree using tools such as iTOL (Letunic and Bork 2016) or GraPhlAn (Asnicar et al. 2015). Alternatively, several online bioinformatics databases offer precomputed summaries of taxonomic distributions for genes based on Linnean taxonomic classification or the NCBI taxonomy (Yang and Bourne 2009; Sayers et al. 2009; Finn et al. 2016; Adebali and Zhulin 2017). However, there is a need for tools that allow users to explore gene/function distributions across a taxonomically curated and highly resolved tree of life.

Here, we present AnnoTree (<u>annotree.uwaterloo.ca</u>), a functionally annotated bacterial tree of life that enables interactive exploration of gene/function annotations across nearly 24,000 bacterial genomes. The phylogeny and taxonomic nomenclature used within AnnoTree is derived from the recently developed Genome Taxonomy Database (GTDB) (Parks et al. 2018). The GTDB overcomes several challenges with the construction of an annotated tree of life as it is *standardized* (its taxonomic nomenclature and phylogeny are made to be internally consistent) and *thorough* (it includes a large number of novel bacterial genomes derived from metagenomic sources). This differentiates the GTDB taxonomy and AnnoTree from similar approaches that rely on the NCBI taxonomy (Federhen 2012), whose hierarchy disagrees with several recent reconstructions of bacterial phylogeny (Bromberg et al. 2016; Hug et al. 2016).

## Results

To construct the AnnoTree database, we re-annotated all 23,936 genomes in the GTDB (Release 02-RS83) using a consistent annotation pipeline. Following gene prediction, we assigned functional annotations [Pfam protein families (Finn et al. 2016) and KEGG orthology (KO) identifiers (Kanehisa et al. 2017)] to protein sequences using standard confidence score thresholds, resulting in 39,153,531 Pfam and 37,850,864 KEGG annotations. All taxonomic information, protein sequences, and functional annotations are stored in a back-end MySQL database for rapid retrieval by the front-end AnnoTree application (**Fig. 1**). To enable phylogenetic visualization of all 23,936 bacterial genomes, AnnoTree divides the bacterial tree of life into distinct views by each major taxonomic level. A user can explore the phylogenetic distribution of a trait anywhere from the phylum to genome level.

**Figure 1.**
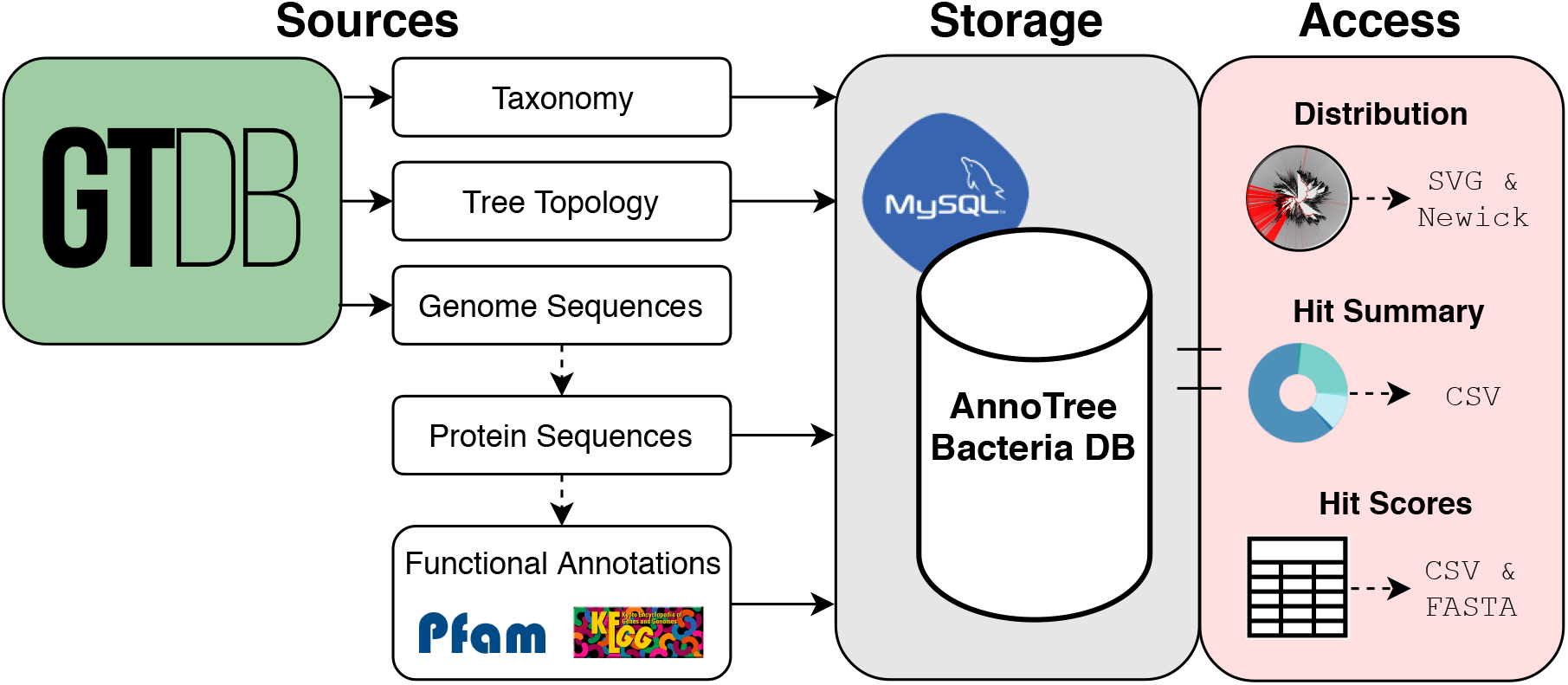
Data flow in the AnnoTree application. Raw values and computed features derived from data obtained from the GTDB is stored in a MySQL database that will be updated to match revisions made to the GTDB. Users can access data relevant to their queries in the form of figures and tables that are rendered in their browser. The figures themselves and the data used to generate them can be downloaded in various file formats from the AnnoTree interface.

AnnoTree can be queried in several ways: by Pfam protein family, KO term, or taxonomic name/id. Additionally, species that appear in a BLAST result can be visualized by uploading the BLAST XML2 output file directly. AnnoTree will then generate a “painted” phylogeny using root-to-tip coloring for all lineages containing matches to the query (**Fig. 2**). Visualizations are also accompanied by basic taxonomic information and distribution summary statistics based on GTDB nomenclature (**Fig. 2**). Publication-quality SVG images, Newick formatted phylogenies for any selected subset of the tree, and taxonomic distribution tables of all queries can be downloaded for offline analysis or editing. Confidence scores (*E*-values) and options for downloading protein sequences for each annotation in a genome or lineage are displayed within a pop-up window when a colored node is selected on the tree.

**Figure 2.**
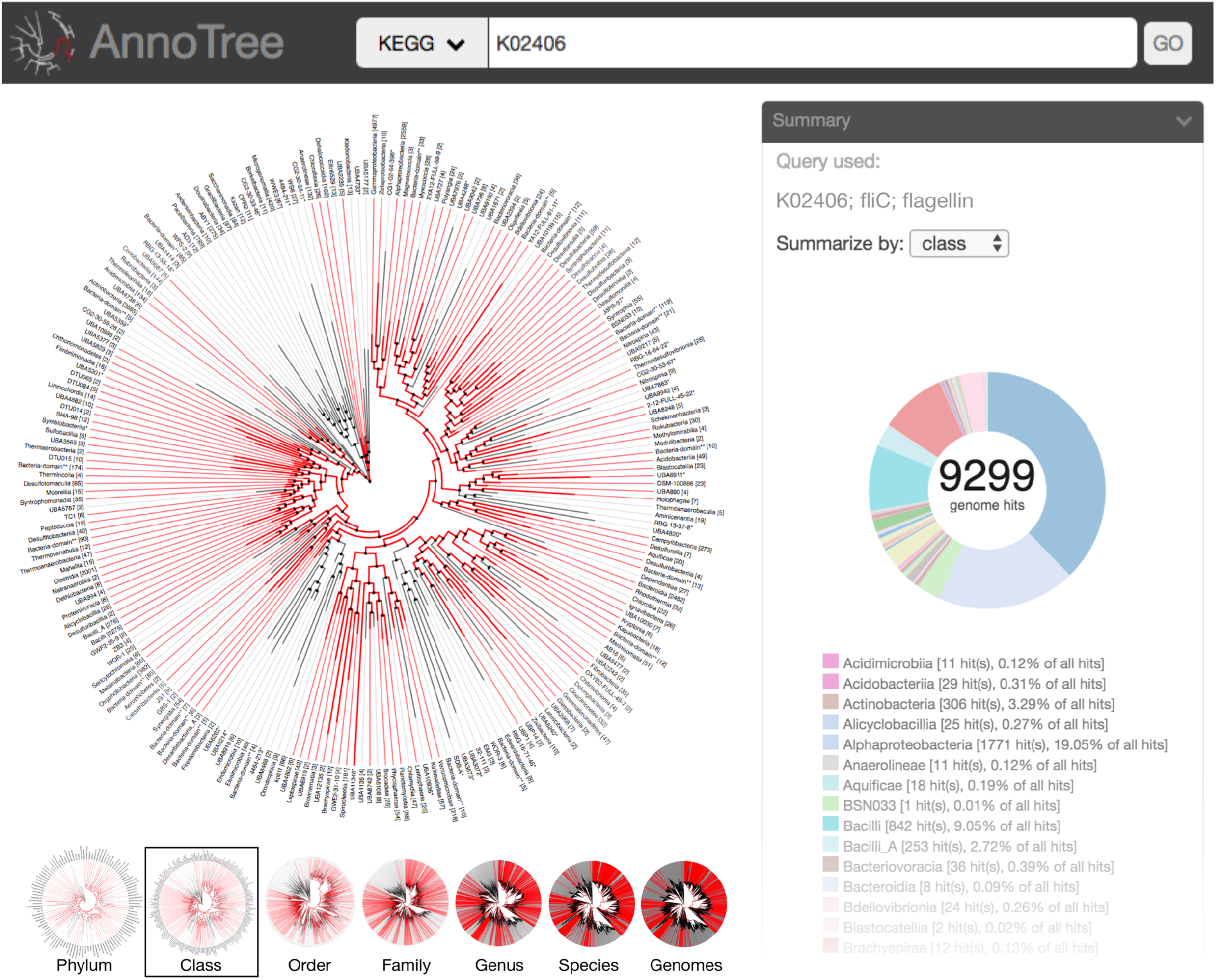
AnnoTree interface overview. AnnoTree can be queried with any number of KO identifiers, Pfam protein families, or NCBI taxon identification numbers to display a mapping of those traits on the GTDB tree at any resolution. Lineages containing at least one genome with the query annotation(s) are highlighted in red. A circle chart displays a taxonomic summary of the genomes containing the flagellin gene (KO identifier: K02406) at a chosen taxonomic level. Smaller trees below show the interactive view when different taxonomic levels are selected by the user. When a highlighted node is clicked, a window appears (not shown in figure) displaying basic taxonomic information, zooming options, and annotation confidence scores.

Since all data is precomputed, users can explore the phylogenomic distribution of any combination of gene families within seconds. As an example, the recent metagenomics-driven discovery of commamox bacteria (van Kessel et al. 2015; Daims et al. 2015) can be reproduced through a simple AnnoTree query by searching for genomes possessing all three key genes that act as a signature for commamox activity: KO terms K00371 (*nxrB)*, K10944 (*amoA)*, and K10535 (*hao*). Highlighted in the tree are the known commamox species (i.e., organisms within the genus *Nitrospira*), along with several additional taxa implicated as having potential commamox-like activity (e.g., *Crenothrix*) (**Supplemental Fig. 1**).

As a second example, the recent discoveries of homologs of important bacterial toxins outside of their respective bacterial lineages can be reproduced and visualized phylogenetically using simple AnnoTree queries. A query with Pfam PF01742 (botulinum neurotoxin protease) reveals a taxonomic distribution outside of *Clostridium* including the lineages *Weissella* and *Chryseobacterium*, consistent with earlier analyses (Mansfield et al. 2015, 2017) (**Supplemental Fig. 2**). Similarly, a search with the diphtheria toxin domains (PF02763 or PF02764) reveals homologs in related genera *Streptomyces* and *Austwickia*, again reproducing recent analyses (Mansfield et al. 2018) almost instantaneously (**Supplemental Fig. 3**). These examples illustrate the use of AnnoTree as a hypothesis-generating tool by revealing distributions of gene families that may be new or unexpected to users.

### Lineage-specific gene families

As an initial exploration of the data within AnnoTree, we examined the distributions of all 77,004,395 Pfam and KO annotations when mapped onto the GTDB bacterial tree of life. Based on the phylogenetic conservation score (τ_D_) (Martiny et al. 2013), 68.1% of KO identifiers and 60.0% of Pfam protein families had significantly non-random phylogenomic distributions (*P* < 0.05), revealing a greater phylogenetic congruency for KO predictions than Pfam predictions. Next, we analyzed the distributions of Pfam and KO annotations to identify those with strong lineage-specificity, which we defined as those where 95% of the members of a gene family occur in 95% of the taxa under a specific phylogenetic node. Based on this criteria, we identified 358 (3.2%) Pfam protein families and 152 (0.9%) KO identifiers with lineage-specific distributions in Bacteria (**Supplemental Data File 1**). We observed a trend in which lineage-specific KO identifiers and Pfam protein families increase in frequency from higher (e.g., phylum) to lower (e.g., species) taxonomic levels (**Supplemental Fig. 4**), consistent with the idea that gene family taxonomic distributions tend to diversify over time and that HGT impacts evolution over short evolutionary timescales (McDonald and Curriea 2017). Although lineage-specific families are relatively rare at high taxonomic levels, these cases often represent ancient, clade-defining bacterial innovations. Examples include K18955 (WhiB family transcriptional regulator) in the Actinobacteria, PF07542 (ATP12 chaperone) in the Alphaproteobacteria, and numerous photosynthesis-related genes within the Cyanobacteria (class *Oxyphotobacteria*).

Lineage-specific gene families can provide insights into the unique biology of their respective organisms. For example, eight lineage-specific Pfam and KO annotations were detected within the *Endozoicomonas* subtree, a clade of endosymbiotic bacteria that inhabit numerous marine eukaryotic hosts (Neave et al. 2016). Consistent with possible utilization of host processes, the lineage-specific genes detected within this clade appear to be of eukaryotic origin and include genes involved in cytoskeletal organization (PF01302), eukaryotic cell-cell signaling (PF00812), apoptosis inhibition (K010343, K010344, K04725, PF07525), and eukaryotic proteolysis (K01378). Given the occurrence of numerous lineage-specific gene families in *Endozoicomonas*, we asked whether lineage-specific gene families may be overrepresented in certain taxa or branches of the bacterial tree. Indeed, lineage-specific genes were significantly enriched in specific taxonomic groups. Notable examples include 37 Pfam protein families within the *Bacillus_A* genus, and 19 Pfam protein families within the Actinobacteria that are largely composed of proteins of unknown function. We also observed an overrepresentation of lineage-specific gene families in numerous well-studied pathogens (e.g., *Bordetella*, *Helicobacter*, *Legionella*, and *Vibrio*) (**Supplemental Figs. 5-7**; **Supplemental Data File 1**). This is in part due to the presence of lineage-specific virulence factors and toxins, but is also likely influenced by annotation bias towards organisms of biomedical interest (Haynes et al. 2018).

### Gene families with patchy distributions

Although 60-68% of functional annotations show a significant phylogenetic signal when mapped onto the tree, more surprising are the remaining 30-40% that show more random phylogenetic distributions, potentially reflecting the widespread horizontal transfer and/or frequent gene gain/loss that is known to occur in bacterial genomes (Ochman et al. 2000; Eisen 2000). To investigate this further, we ranked all Pfam and KEGG annotations according to their phylogenetic patchiness, determined by homoplasy score (total number of gains and losses by parsimony) normalized by gene family size after filtering out traits with family size less than 50 (**Supplemental Data File 2**, see Methods). KEGG KO terms were grouped into their higher-level functional categories for visual comparison of broader trends (**Fig. 3, Supplemental Data File 3**). Not surprisingly, “viral” (bacteriophage) genes ranked the highest in homoplasy in both Pfam and KEGG annotations, and therefore are the single most phylogenetically scattered class of genes in bacteria. In contrast, gene functions with extremely low homoplasy include sporulation, photosynthesis, and core processes such as transcription, replication, and protein synthesis (**Fig. 3**). Highly scattered genes showed significant overrepresentation among specific taxonomic groups such as the genera *Pseudomonas_E, Streptomyces,* and *Mycobacterium* (**Supplemental Data Files 4, 5**), suggesting that these taxa may be taxonomic “hotspots” of HGT.

**Figure 3.**
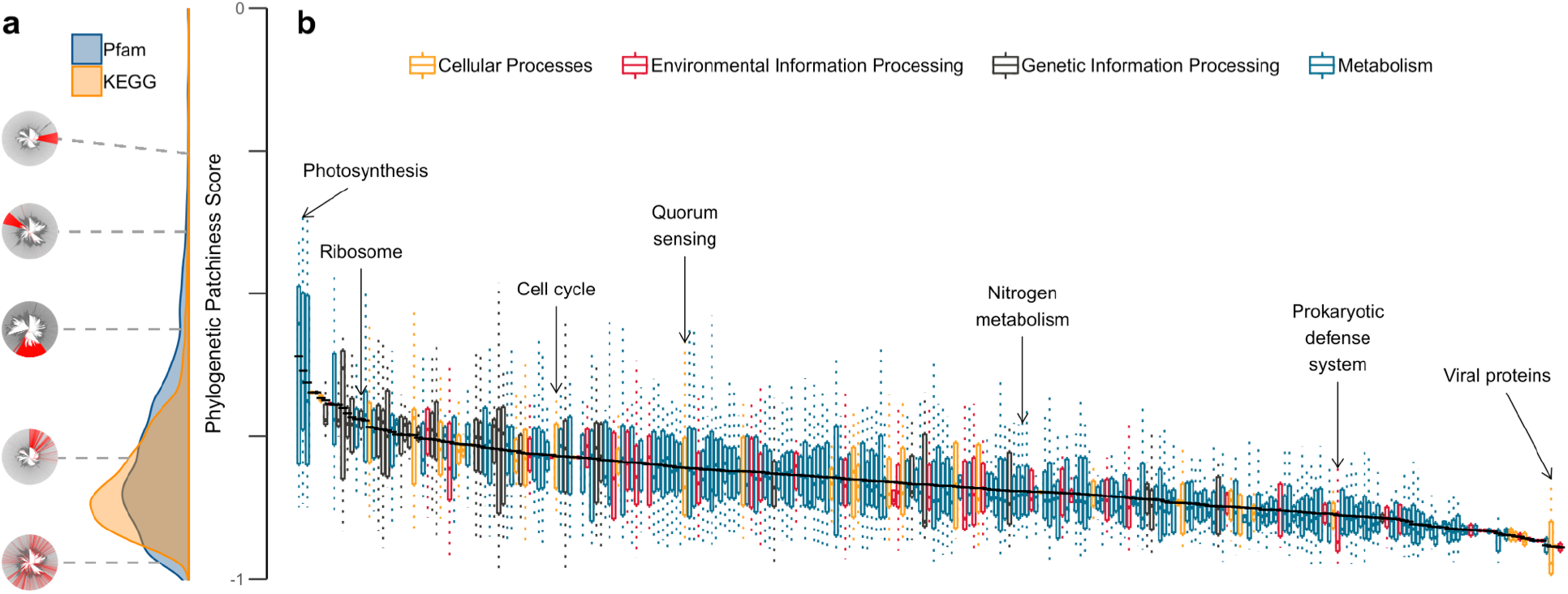
Phylogenetic patchiness of annotations inferred using AnnoTree. Phylogenetic patchiness was computed for each KEGG KO identifier and Pfam protein family using the consistency index (CI), a common homoplasy metric representing the inverse of the minimum possible number of state changes (trait gain or loss) given the tree topology. The final phylogenetic patchiness score is equal to log(CI)/log(family size) where family size is the total number of genomes containing the trait. (A) Density plot showing the distribution of phylogenetic patchiness scores of Pfam protein families and KO identifiers with different visual examples of varying patchiness (red=present; gray=absent). The phylogenetic distribution plots are, from top to bottom: K10922 (transmembrane regulatory protein ToxS), K18955 (WhiB transcriptional regulator), PF01848 (ATP12 chaperone), PF01848 (Hok/Sok antitoxin system), and K07495 (putative transposase). (B) Mean-sorted box plots containing phylogenetic patchiness scores of KO identifiers in their respective KEGG pathways and KEGG BRITE categories. The mean patchiness score of a set of KO identifiers in a KEGG pathway or KEGG BRITE category is indicated by a black line.

We then examined in more detail the top 100 gene families that showed the most scattered distributions across the bacterial tree. Not surprisingly, this list of gene families is dominated by transposases, CRISPR- and bacteriophage-associated gene families (**Supplemental Data File 2**). Numerous gene families of unknown function were included among the most patchy gene families, but further examination revealed that most of these genes are likely bacteriophage-derived. The extreme phylogenetic patchiness of bacteriophage and CRISPR genes is not only consistent with their known evolutionary dynamics but could also reflect the ongoing “arms race” between these two opposing biological forces (phage infection versus phage defense). Other biologically relevant members of the 1% most highly scattered KO genes include: K19057-K19059 (*merC*, *merD*, and *merR* of the *mer* operon) for mercury resistance; K19155 and K19156, components of a toxin-antitoxin system characterized in *E. coli*; K15943, K15945, and K16411 for polyketide antibiotic biosynthesis; and K19173-K19175 for DNA backbone S-modification (phosphorothioation) (**Supplemental Data File 2**).

### Reductive dehalogenases

As a case study for the hypothesis generation and data mining strengths of AnnoTree, we selected a gene family of significant biological interest that ranked among the top percentile of homoplasy scores: *pcpC*; tetrachloro-p-hydroquinone reductive dehalogenase (K15241) **Supplemental Data File 2**). As key enzymes in bioremediation of chlorinated solvents, there has been extensive characterization of the diversity and phylogenomic distribution of reductive dehalogenases (Rdhs) and organohalide respiring organisms (Hug et al. 2013). Using AnnoTree, we compiled a dataset of Rdh genes and associated taxa using Pfam query PF13486. Our analysis produced a comprehensive dataset of 1,299 putative Rdh genes from 385 genera and 38 phyla (**Supplemental Table 1**, **Supplemental Figs. 8, 9**), which not only recapitulates the known diversity of Rdh-associated phyla, but significantly expands it. In comparison, a manually-curated Rdh-specific database contains 264 Rdh genes from only 19 genera and 6 phyla (Hug et al. 2013), less than 15% of the total diversity identified by AnnoTree (**Supplemental Table 1**). The AnnoTree-derived dataset includes several newly predicted *rdh*- encoding taxa discovered from metagenome-assembled genomes (**Supplemental Table 2)**, including the candidate phyla KSB1 (4 of 6 genomes, *rdh* copy number = 1) and UBP10 (7 of 14 genomes, *rdh* copy number = 1), as well as *Rhodospirillales* UBA2165 (*rdh* copy number = 13) and *Acidobacterium* UBA2161 (*rdh* copy number = 8) (**Supplemental Fig. 9**, **Supplemental Table 2**). The novel organisms with high *rdh* copy numbers are potential obligate organohalide respirers and may be valuable for remediation efforts. By revealing both known and potentially novel groups of organohalide respiring bacteria, the Rdh case study highlights the ability of AnnoTree to capture a broad and complete taxonomic diversity of a gene family, with accompanying hypothesis generation around the evolution and ecology of a function of interest.

## Discussion

Ultimately, by combining functional annotation data with evolutionary data, AnnoTree provides an automated framework for users to explore the distribution of function across the bacterial tree of life. These visualizations allow users to investigate a wide variety of research questions concerning their genes and functions of interest. As starting points for future analyses, we have assessed Pfam and KO annotations based on phylogenetic conservation, homoplasy, and lineage-specificity. However, while AnnoTree provides a snapshot of gene occurrence, additional sequence and phylogenetic analyses are required to validate many of these predictions. Our development team has plans to include additional functional annotation types and to provide a standalone version of AnnoTree for local use. The AnnoTree database will also be continuously and automatically updated to reflect revisions of the GTDB taxonomy as the data become available. Future work will expand AnnoTree’s taxonomic scope to Archaea and Eukaryotes. We anticipate that AnnoTree will become a valuable resource for exploring the evolution and phylogenomic distribution of genes and functional traits across the tree of life.

## Methods

### Gene prediction, annotation, and profile generation

Gene prediction was performed with Prodigal v2.6.3 (Hyatt et al. 2010) and annotated using the Pfam v27 (Finn et al. 2016) and UniRef100 (Suzek et al. 2015) (downloaded March 6, 2018) databases. Pfam protein families were identified using HMMER v3.1b1 (Eddy 2011) with model specific cutoff values for the Pfam (-cut_gc) HMMs. Pfam annotations were assigned using the same methodology as the Sanger Institute, which accounts for homologous relationships between Pfam clans (see pfam_scan.pl on the Sanger Institute FTP site). UniRef100 was used to establish KO annotations by creating a DIAMOND v0.9.22 (Buchfink et al. 2015) database consisting of all UniRef100 clusters with one or more KO identifiers. KO identifiers were then assigned to predicted genes through homology with the following criteria: *E*-value cutoff ≤1e-5, percent identity ≥30%, and query and subject percent alignments ≥70%. A count matrix was computed for each trait and genome combination based on the annotation methods described above. The count matrices were converted to binary presence/absence profiles for all analyses, where a genome with at least one qualifying hit score for a trait was assigned ‘1’ and ‘0’ otherwise.

### Web application development

AnnoTree has three components: a front-end, back-end, and a MySQL database. GTDB (Parks et al. 2018) (Release 02-RS83) provides bacterial taxonomy structure data, along with the presence and absence of Pfam domains and KEGG genes in each genome assembly. Those are imported to MySQL tables. The back-end is a Python Flask application to serve REST API endpoints. It converts JSON query to SQL statements. The front-end is a single page application using modern web frameworks such as D3, React, and Mobx. The tree and summary chart is drawn using D3.js, while other UI components are encapsulated by React. Mobx is a state management engine that triggers UI update whenever state variables change.

### Calculation of phylogenetic conservation

The trait depth (τ_D_) for each annotation profile on the GTDB tree was calculated using the consenTRAIT algorithm (Martiny et al. 2013) implemented in the castor R package (Louca and Doebeli 2018). A trait was classified as phylogenetically conserved if the probability of encountering a profile with such a τ_D_ or higher is less than 5% (ie. *P* < 0.05) based on 1000 different independently- and randomly-drawn binary presence/absence profiles where the probability of a tip exhibiting the trait is equal to the proportion of positive states in the trait’s profile.

### Classification of lineage-specific traits

Traits were classified as lineage-specific if there was at least one clade in the tree where at least 95% of presence states occurred in at least 95% of the taxa in that clade and that no more than half of the genomes in the tree contained the trait. The node furthest from the root of the GTDB tree passing these criteria was assigned the root of the lineage-specific clade for that trait. The trait’s taxonomic rank was selected as the lowest identical taxonomic rank between all genomes of the lineage-specific clade.

### Calculation of homoplasy metrics

Parsimony-based homoplasy metrics were used to quantify phylogenetic scatter of traits. The consistency index (CI) and retention index (RI) were calculated for each annotation profile with the GTDB tree using the phangorn R package (Schliep 2011). The homoplasy slope ratio (HSR) was calculated similarly with a custom script (“HSR.R” in https://bitbucket.org/doxeylab/annotree-scripts) that utilizes the algorithm described in Meier *et al*. (1991). The random homoplasy slope was calculated using 100 randomly-drawn presence/absence profiles with equal probability of presence and absence.

### Taxonomic rank homoplasy enrichment analysis

Annotations contained within less than 50 genomes were removed before verifying taxonomic enrichment of homoplasic traits for each annotation type. Taxonomic rank presence/absence profiles for each trait were generated for each taxonomic rank by combining the profiles of all encompassing genomes; ‘1’ was assigned if at least one genome possessed the trait and ‘0’ otherwise. Next, traits were ranked by increasing log[CI]/log[family_size]. Each taxonomic rank at each taxonomic level was tested for over-enrichment within the 5% most homoplasic traits in Bacteria (KO: 618; Pfam: 552) using the hypergeometric test. The tests were conducted similarly to those done by Nasir *et al*. (2012). *P* values were obtained using the fisher.test function of R with the ‘alternative’ option set to ‘greater’.

The contingency table was given as follows:

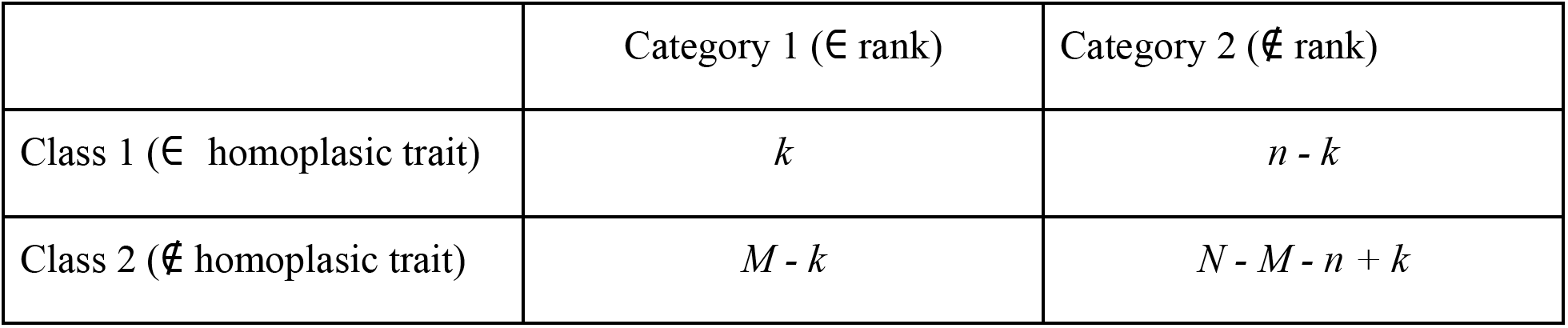

where *k* is the number of unique homoplasic traits within the rank, *n* is the number of unique ranks that contain at least one of the homoplasic traits, *M* is the total number of unique traits within the rank, and *N* is the total number of unique traits. *P* values were corrected for multiple tests at each taxonomic level using the Benjamini-Hochberg method (Benjamini and Hochberg 1995).

### Data Availability

The AnnoTree application is available at http://annotree.uwaterloo.ca. All software and data used within AnnoTree can be downloaded at: http://annotree.uwaterloo.ca/downloads.html. Additional data for the genomes and taxonomy derived from the GTDB can be found at: http://gtdb.ecogenomic.org/downloads.

## Acknowledgements

This work was supported by an NSERC Discovery Grant and Ontario Early Researcher award to A.C.D. DHP was supported by the Australian Research Council Laureate Fellowship (FL150100038) awarded to Philip Hugenholtz (P.H). LAH is supported by a Tier II Canada Research Chair. We also thank Lee Bergstrand for technical help and Philip Hugenholtz for helpful suggestions.

## Author contributions

H.C and K.M. built the front-end interface of AnnoTree and back-end database. K.M. performed data analysis. D.H.P. assisted with bioinformatic analysis and genome annotation. L.A.H. assisted with phylogenetics and tool design. A.C.D. and K.M. wrote the manuscript. A.C.D. conceived the project and tool design, and supervised H.C. and K.M.

